# A physicochemical roadmap of yeast replicative aging

**DOI:** 10.1101/858720

**Authors:** Sara N. Mouton, David J. Thaller, Matthew M. Crane, Irina L. Rempel, Anton Steen, Matt Kaeberlein, C. Patrick Lusk, Arnold J. Boersma, Liesbeth M. Veenhoff

## Abstract

Cellular aging is a multifactorial process that is characterized by a decline in homeostatic capacity, best described at the molecular level. Physicochemical properties such as pH and macromolecular crowding, are essential to all molecular processes in cells and require maintenance. Whether a drift in physicochemical properties contributes to the overall decline of homeostasis in aging is not known. Here we show that the cytosol of yeast cells acidifies modestly in early aging and sharply after senescence. Using a macromolecular crowding sensor optimized for long-term FRET measurements, we show the macromolecular crowding changes less in longer-lived cells in contrast to shorter-lived cells. While the average pH and crowding levels change only modestly with aging, we observe drastic changes in organellar volume, leading to crowding on the µm scale, which we term organellar crowding. Our measurements provide an initial framework of physicochemical parameters of replicatively-aged yeast cells.

## Introduction

Cellular aging is a process of progressive decline in homeostatic capacity ^1, 2^. Key molecules govern the aging process and can extend health and lifespan by maintaining prolonged homeostasis ^3^. The function of these molecules ultimately depends on physicochemical properties, such as pH, macromolecular crowding, and ionic strength, all of which require maintenance. For example, the formation of membrane-free compartments is particularly sensitive to crowding and pH levels. Thus, age-associated changes in macromolecular crowding or pH have the potential to drive profound changes to intracellular organization ^4–6^. However, we do not have a clear understanding of the physicochemical parameters in cells or how they change with age. *Saccharomyces cerevisiae* is an excellent model system to determine these parameters because single-cells can be directly monitored by microscopy as they age ^7^. Importantly, many of the molecular mechanisms that contribute to yeast aging are conserved in humans ^8^.

pH is a crucial physicochemical parameter in aging because it affects most, if not all, biochemical processes. The main contributors to pH homeostasis in the yeast cytoplasm are a P-type ATPase, Pma1, and the V-ATPase. Pma1 localizes in the plasma membrane and transports cytosolic protons out of the cell ^9–11^, while the V-ATPase pumps protons from the cytosol into the lumen of various organelles and regulates their pH. Pma1 is asymmetrically retained in the mother cell and during cell divisions its levels increase in replicative aging, resulting in the alkalinization of the mother’s aging cell cortex (region close to the plasma membrane) ^12^. Further, the aging yeast vacuoles lose acidity early in life, which has adverse downstream effects on mitochondrial functionality and potentially also on DNA repair ^13^. This change in vacuolar pH homeostasis could be induced, at least in part, by the V-ATPase that loses protein complex stoichiometry with aging, potentially reducing the number of functional complexes ^14^. This relation is further shown by partial recovery of the stoichiometry, after overexpression of the Vma1 component, which subsequently increases lifespan in yeast ^13, 14^. The importance of pH homeostasis in aging is also apparent in humans. Human senescent cells show increased lysosomal content and increased lysosomal pH, and in age-related pathologies such as Alzheimer’s and Parkinson’s disease, lysosomes are dysfunctional ^15^. V-ATPases are highly conserved, and Pma1 shares structural and functional similarities with the Na^+^K^+^ ATPases in mammalian cells ^16–18^. Hence, pH needs to be considered when defining a physicochemical roadmap of cellular aging.

pH also influences macromolecular organization in yeast: Starvation acidifies the cytoplasm, which leads to a phase transition to a “gel-like” state that hampers diffusion of µm-sized particles ^4^. The propensity to undergo a phase transition is amplified by macromolecular crowding ^19^. Cells are highly crowded with macromolecular concentrations estimated to be between 80 to 400 mg/ml ^20, 21^. Macromolecular crowding retards diffusion, influences protein volume and association equilibria ^22–24^, including e.g., condensate formation *in vitro* and *in vivo* ^5, 25^. These effects are caused by steric exclusion, next to weak chemical interactions ^26–28^, and depend on the concentration, size, and shape of the molecules involved and are larger when crowders are smaller-sized than the reacting molecule ^26, 29^. For example, an increased number of ribosomes slows down diffusion of 20 nm and 40 nm particles, but not protein-sized molecules ^5^.

It is not trivial to quantitatively measure crowding in living organisms ^26^, and most estimates of crowding were obtained from measuring dry cell mass ^20, 21^, which does not reflect the actual crowding. We previously developed a genetically encoded FRET-based probe that allows quantification of macromolecular crowding *in vivo* ^30^. This probe was further optimized to be applied in bacteria and yeast ^31^ and allows a spatiotemporal readout of changes in crowding ^30, 32^. As it is non-invasive, this probe is ideal to determine changes in macromolecular crowding during yeast replicative aging.

The volume of a cell needs to be coupled to biopolymer synthesis to maintain macromolecular crowding ^33, 34^. For example, it was suggested that oversized yeast cells have reduced molecular density ^35^. One of the most dramatic features of yeast is a marked increase in cell size ^36–38^. Concomitantly, yeast organelles like vacuoles ^39^, the nucleus (including nucleoli ^40, 41^), and mitochondria ^42^ can exhibit changes in morphology. These changes in compartment size and shape directly impact the cytosolic volume while presenting physical barriers to molecular movement and surfaces on which molecules can be adsorbed. Furthermore, changes in compartmental volume alters energy consumption. For example, a small compartment, such as the endosome, needs to import fewer protons to maintain pH compared to larger compartments ^43^. Additionally, specific cell types have different organelle sizes related to their function: for example, secretory cells have expanded ER ^44^. In healthy young cells, a “volume hierarchy” is observed where the cytosol represents the largest compartment, followed by the nucleus and vacuole. However, despite its importance, it is not clear how, or whether, these compartmental volumes change during aging.

Here we present a framework describing how critical physicochemical parameters change during yeast replicative aging. We find that the cytosol shows a modest acidification in early aging but drops more significantly after the cells stop dividing. We optimize our macromolecular crowding sensors for long-term FRET measurements and show that longer-lived cells tend to maintain macromolecular crowding better than shorter-lived cells. While the pH and crowding change only modestly in aging, we observe drastic changes in organellar volumes, leading to crowding on the µm scale, which we term organellar crowding.

## Results

### Yeast replicative aging leads to acidification of the cytosol, especially after entry into senescence

To follow cytosolic pH levels in aging, we utilized the fluorescence based, genetically encoded pH sensor ratiometric pHluorin ^45^. pHluorin is a GFP variant that responds to changing pH: With increasing acidity, the excitation at 395 nm decreases, while the excitation at 475 nm increases. We expressed pHluorin in BY4741 background strain and recorded F390/F475 ratios from aging cells using the ALCATRAS microfluidic device ^7^. We followed single-cell life histories from 30 cells over 80 h by wide-field fluorescence microscopy (Fig. 1*A*). We have performed an *in vivo* calibration of pHluorin (Fig. 1*B*) and found that young cells have an average cytosolic pH of 7.7 and that pH levels are comparable between individual cells (Fig. 1*C*). The measured average cytosolic pH is higher than the previously reported pH of 7.3 ^46^. The higher value could be because cells trapped in the microfluidic devices experience stable pH and nutrient availability, which is not the case when imaging cells derived from a culture on a coverslip, where nutrients are limiting.

**Figure 1.**
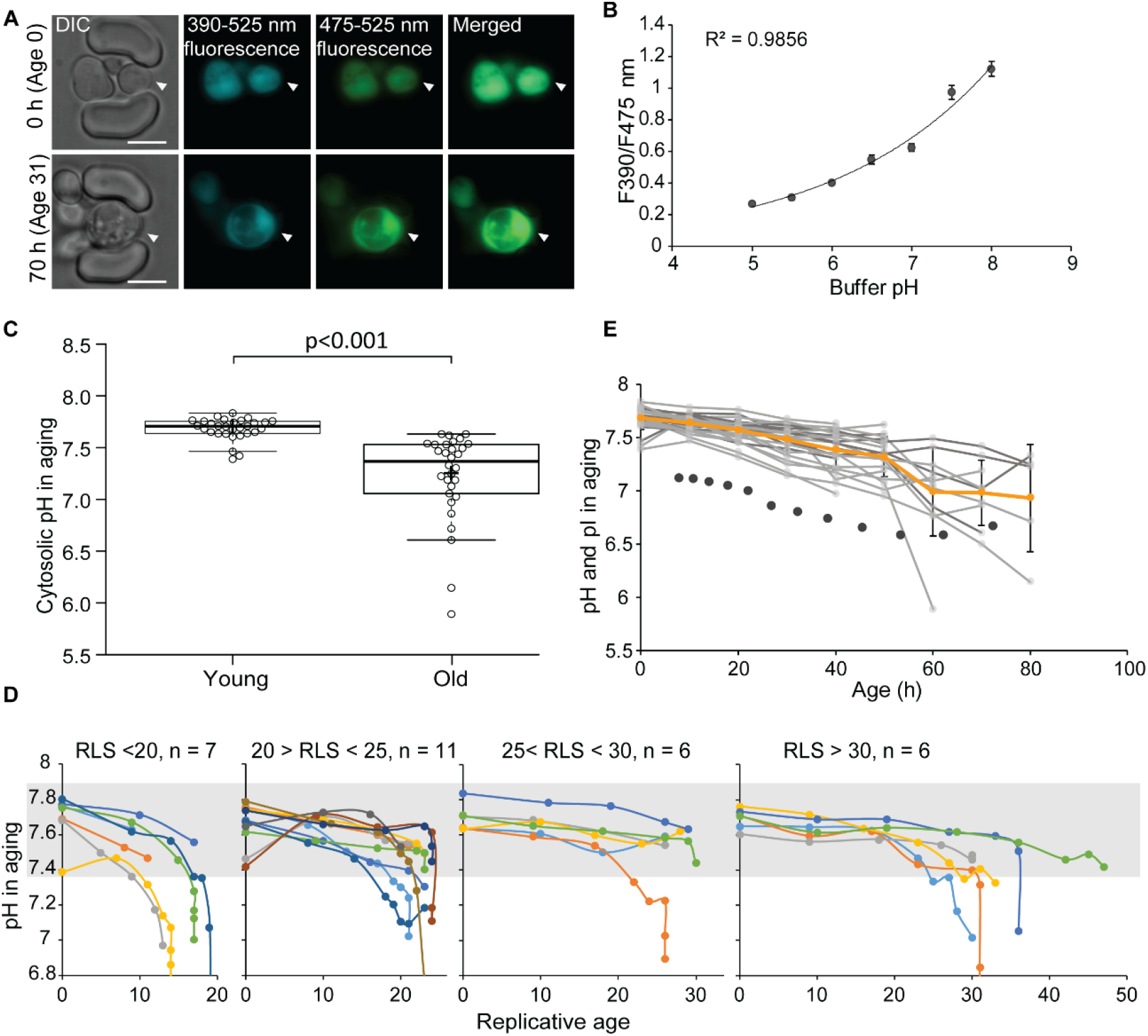
The cytosol of old cells acidifies, and cells display more substantial variability in cytosolic pH with aging. (*A*) Representative images of cells expressing ratiometric pHluorin imaged at the start of the experiment and after 80 hr; the replicative age is indicated. Young cells are trapped in the microfluidic device, and bright-field images are taken every 20 min to determine the age of the cells. Fluorescent images are taken once every 10 h with excitation at 390 and 475 nm and emission at 525 nm. The images show DIC, DAPI, and FITC channels and merged fluorescence. The scale bar represents 5 μm. The trapped aging cell is indicated with white arrowheads. B) Calibration curve showing the relationship between intracellular pH and pHluorin ratios. Cells expressing pHluorin from an exponential culture at OD_600_ of 0.5 were resuspended in buffers, titrated to pH 5, 5.5, 6, 6.5, 7, 7.5, 8, containing final concentrations of 75 μM monensin, 10 μM nigericin, 10 mM 2-deoxyglucose, and 10 mM NaN_3_. Monensin and nigericin are ionophores that carry protons across the plasma membrane, while the 2-deoxyglucose and NaN_3_ deprive the cell from ATP and thus block energy-dependent pH maintenance. Each point represents data from 20 cells. (C) Data collected from 30 yeast cells during the process of replicative aging. The young group consists of data points from the first recorded ratio at time 0 h, and the old group consists of the last recorded measurement before cell death. Crosses indicate the average and bold lines show the median, p = 4.9E-07. (D) Single-cell profiles of the pH at different replicative ages. Grey bar indicates the range of pH values measured in the first time point (young age) to illustrate that around half of the cells remain within this range throughout their lifespan. (E) The pH of single cells during aging plotted versus their age in hours. Each grey line represents a single cell. The orange line shows the average pH. The dark grey circles show the change in total proteome pI in the aging of 1229 proteins. Error bars represent the standard deviation.

To directly compare differences in pH between young and aged cells, we grouped the first and the last measurements taken for each cell under two categories: “young” and “old”. The young category contains cells of predominantly age 0, since new-born daughters are the most common group in an exponentially growing culture. The “old” category is composed of senescent and non-senescent cells of different replicative ages, where most cells had no divisions left to complete (n = 22, 100% of RLS), and others had up to 4 divisions to complete (n = 8, 4-25% of RLS). The average lifespan of this cohort is 25 divisions.

Our results show that the average cytosolic pH significantly decreases by 0.5 pH units in old cells, compared to young (Fig. 1*C*). Furthermore, the pH of individual old cells spans a wide range of pH values from 7.6 to 5.9. Interestingly, 50% of the old cells manage to maintain pH levels above 7.4, which is the lower boundary in young cells, while the remainder of the old cells have a lower pH. The large spread at old ages is related to a substantial drop in pH in cells that stop dividing and enter senescence (Fig. 1D). When we plot the pH of single-cell trajectories for different ranges of replicative life spans, we observe a gradual decrease in pH already early in life in almost all cells, but especially so in those that are short-lived. Indeed, the change in pH in the earlier divisions (up to replicative age 15) correlates to the cell lifespan with Spearman correlation of 0.6 (p = 0.001) (Fig. S1*A*), but we find no relationship between the lifespan of cells and (i) pH at young age, (ii) the pH at old age, or (iii) the change in pH (Fig. S1*B-D*). Supported by previous measurements of pH in the vacuole and at the plasma membrane (cell cortex) in yeast replicative aging ^12, 13^, these findings are indicative of loss of global pH homeostasis as yeast age.

The activity of proton pumps as well as the strong buffering capacity of e.g., metabolites and amino acid side chains determine pH homeostasis. Because the concentration of amino acids with a physiologically relevant pKa at protein surfaces is orders of magnitude larger than the concentration of free protons at pH 7 (protein concentration is in the mM-range while pH 7 corresponds to 60 nM H^+^), the proteome represents a buffer for changes in pH. We thus assessed the proteome isoelectric point (pI) during yeast replicative aging. We utilized available datasets for protein abundance during aging ^14^, predicted isoelectric point (Saccharomyces Genome Database, SGD), and protein copy number ^47^. We used data from 1229 proteins and corrected our calculations for relative protein abundance, thus weighing the proteome pI for the copy number of each protein. We found that in young cells, the proteome pI is 7.1, which accounts mostly for the cytosolic part of the cell, according to the Panther database for gene ontology. In cells aged for 60 hours (average replicative age ∼22 divisions), the proteome pI reaches as low as 6.7 (Fig.1*E*). Interestingly, the proteome pI roughly follows the pH of the cytoplasm during aging. The analysis of the cytosolic pH and proteome pI in aging, together with previous studies on the activity of proton pumps ^12, 13^, suggests that pH homeostasis is disrupted not only at the level of proton pumps but also at the level of the buffering capacity of the proteome.

### Validation of FRET-based crowding probes during yeast aging

Because pH has previously been related to macromolecular organization ^4, 19^, we asked whether macromolecular crowding also changes in aging yeast cells. To measure macromolecular crowding, we utilized a FRET-based probe for quantification of crowding that we previously developed ^30^, named crGE, and harboring mCerulean3 as donor and mCitrine as acceptor ^31^. We also developed a new probe, named CrGE2.3 by exchanging the donor and acceptor for mEGFP and mScarlet-I, respectively (Fig. 2A, left).

**Figure 2.**
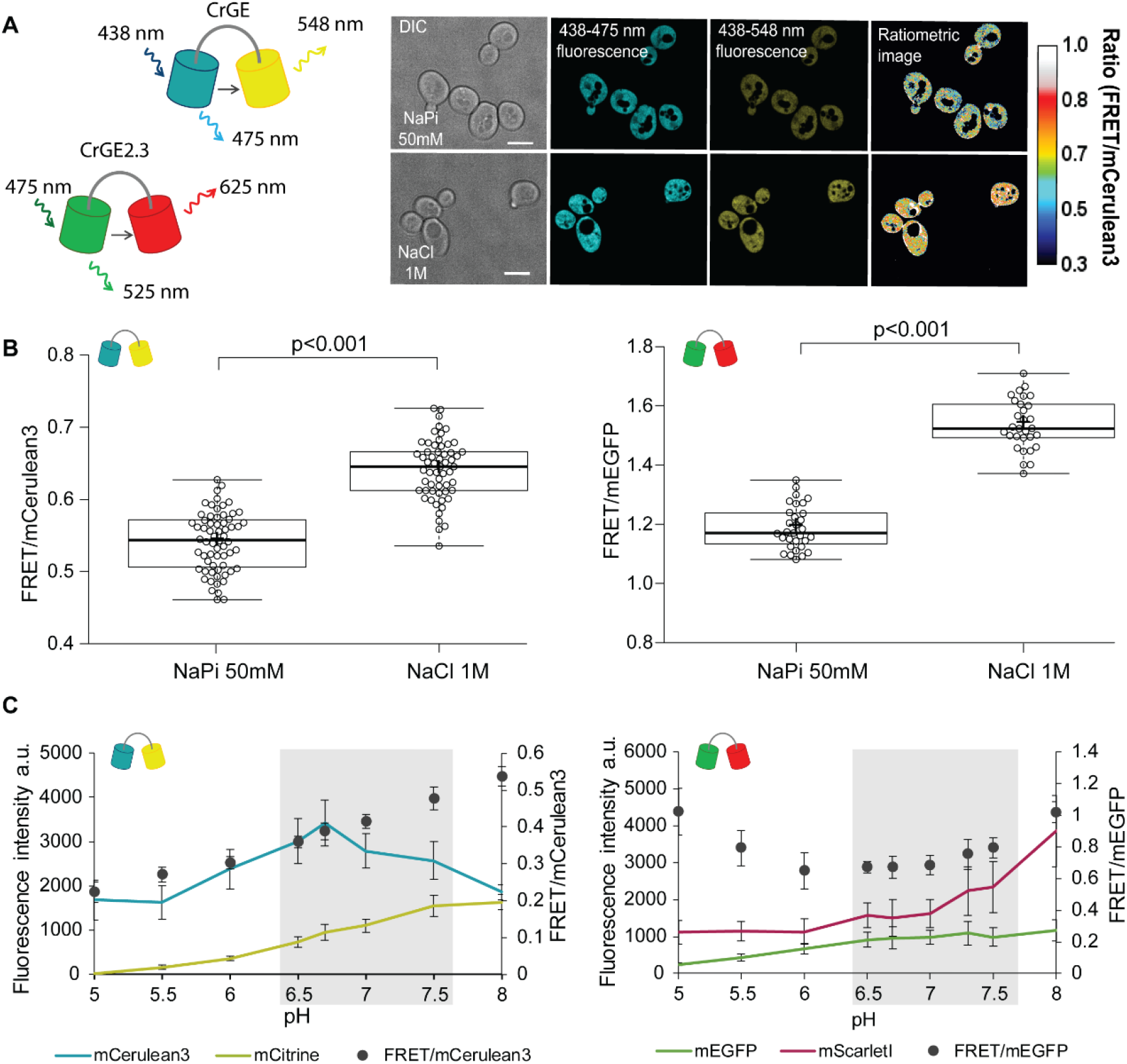
Crowding sensor with traditional CFP-YFP FRET pair is functional in yeast cells, but sensitive to pH variations; the sensor CrGE2.3 with mEGFP-mScarlet-I is aging-compatible. (A) Left: Schematic representation of the crGE and CrGE2.3 sensors. Right: Yeast cells expressing crGE sensor, the DIC, mCerulean3, and FRET channels are shown. Ratiometric image shows changes in the crowding ratio upon osmotic shock. The scale bar is 5 μm. (B) CrGE sensor (left), expressed under a strong constitutive TEF1 promoter in yeast, shows an increase in FRET/CFP ratio upon osmotic shock with 1M NaCl versus control in sodium phosphate buffer at pH 7. Crosses indicate the average and bold lines show the median. The data is from three independent biological replicates with n>58 cells per condition; p=2.0E-25. Right: CrGE2.3 sensor, expression and osmotic shock experiment is the same as for the original CrGE sensor, the data is from 30 cells per condition, p=1.6E-24 (C) Left: Fluorescence intensities of mCerulean3 (blue line), and mCitrine (yellow line; directly excited) and FRET/CFP ratios (black circles). Right: Fluorescent intensities from mEGFP (green line) and mScarlet-I (magenta; directly excited) and FRET/mEGFP ratio (black circles) from permeabilized cells, expressing the crowding sensor CrGE (left) and CrGE2.3 (right), in buffers with pH ranging from 5 to 8. Each data point represents the average of 20 cells; error bars show standard deviation.

Through fluorescence microscopy, we determined the crowding read-out from the ratio of the intensity in the FRET channel and the donor channel (an example is shown in Fig. 2*A*, *right*). To test the validity of the probes, we induced a hyperosmotic shock by resuspending cells from an exponentially growing culture in 1M NaCl. Upon osmotic upshift, the water content and the cell volume should decrease, resulting in increased macromolecular crowding. Indeed, we observed a significant increase (p<0.001) in the crowding ratio of 18% from 0.54 to 0.64 from three independent experiments for crGE (Fig. 2*B*, *left*). To determine whether in the new sensor, crGE2.3 functionality is retained after exchanging the fluorophores, we again induced hyperosmotic shock with 1M NaCl but also 1.5M Sorbitol and observed significant increases in crowding ratio (p>0.001) of 23% from 1.19 to 1.54 for NaCl and to 1.46 for Sorbitol (Fig. 2*B*, *right* and S2*I*). These results show that both sensors are responsive to crowding changes in yeast in a reproducible manner.

In order to perform crowding measurements in aging cells, the sensor should be insensitive to age-related changes, such as proton concentration or fluorescent protein maturation. Since, the cytosolic pH decreases with age it could compromise read-outs from the crowding sensor. To estimate the pH sensitivity of the crGE and the crGE2.3 probe, we resuspended cells permeabilized with ionophores in buffers with pH ranging from 5 to 8. Our results show that readouts from sensors harboring the original FRET pair, mCerulean3-mCitrine, have a strong linear dependence on the intracellular pH (Fig. 2*C*, *left*), in contrast to purified probe in buffer ^30^. Fluorescent proteins were recently shown to have different pH sensitivity *in vivo* compared to *in vitro* ^48^. When assessing the pH sensitivity of the new sensor, CrGE2.3, we found that this probe gives more stable readouts within the pH range observed in aging (Fig. 2*C*, *right*).

Fluorescent protein maturation also plays a role: Aging cells do not maintain the same division frequency, but transition to a slow and irregular division mode, called the senescence entry point (SEP) (Fig. S2, *top row*) ^38^. When the SEP occurs, the relative rates of sensor synthesis, degradation, and dilution through division are all disrupted, leading to an increase in fluorescence of the slower maturing fluorescent protein (Fig. S2*A*). For ageing studies, the FRET pair in crGE2.3, which has similar pH sensitivity and maturation kinetics between the two fluorescent proteins is thus more suitable than the original sensor. To eliminate additional systematic errors, we corrected for an unequal number of donor and acceptor fluorophores by FRET normalization (N_FRET_) ^49^ as pH-induced fluorescence quenching leads to a different number of fluorescent proteins and the same accounts for the proportion of fully matured sensors. This normalization retains readouts from crowding changes (Fig. S2). Thus, in crGE2.3 we have increased the robustness of the probe to make it suitable for challenging long-term aging experiments.

### Crowding homeostasis is maintained during yeast replicative aging

To determine age-associated changes in macromolecular crowding, we constitutively expressed the optimized crowding probe in yeast cells immobilized in the ALCATRAS microfluidic device ^7^. As with the pH-sensor, we observe that the crowding probe localizes in the cytosol and nucleus, but it is excluded from other membrane enclosed organelles (Fig. 3*A* *left*). We collected data from 51 cells in the time course of 70 h and determined N_FRET_ and FRET/mEGFP ratios (Fig. 3*A* *right*). Our measurements show heterogeneity in crowding levels in young cells and more so in old cells (Fig. S3A). Overall, we find that macromolecular crowding is maintained in the course of aging, with average ratios of 0.51 for both young and old cells, where, as in figure 1, the ‘young’ group reflects the first and the ‘old’ group the last crowding measurements (Fig. 3*B*).

**Figure 3.**
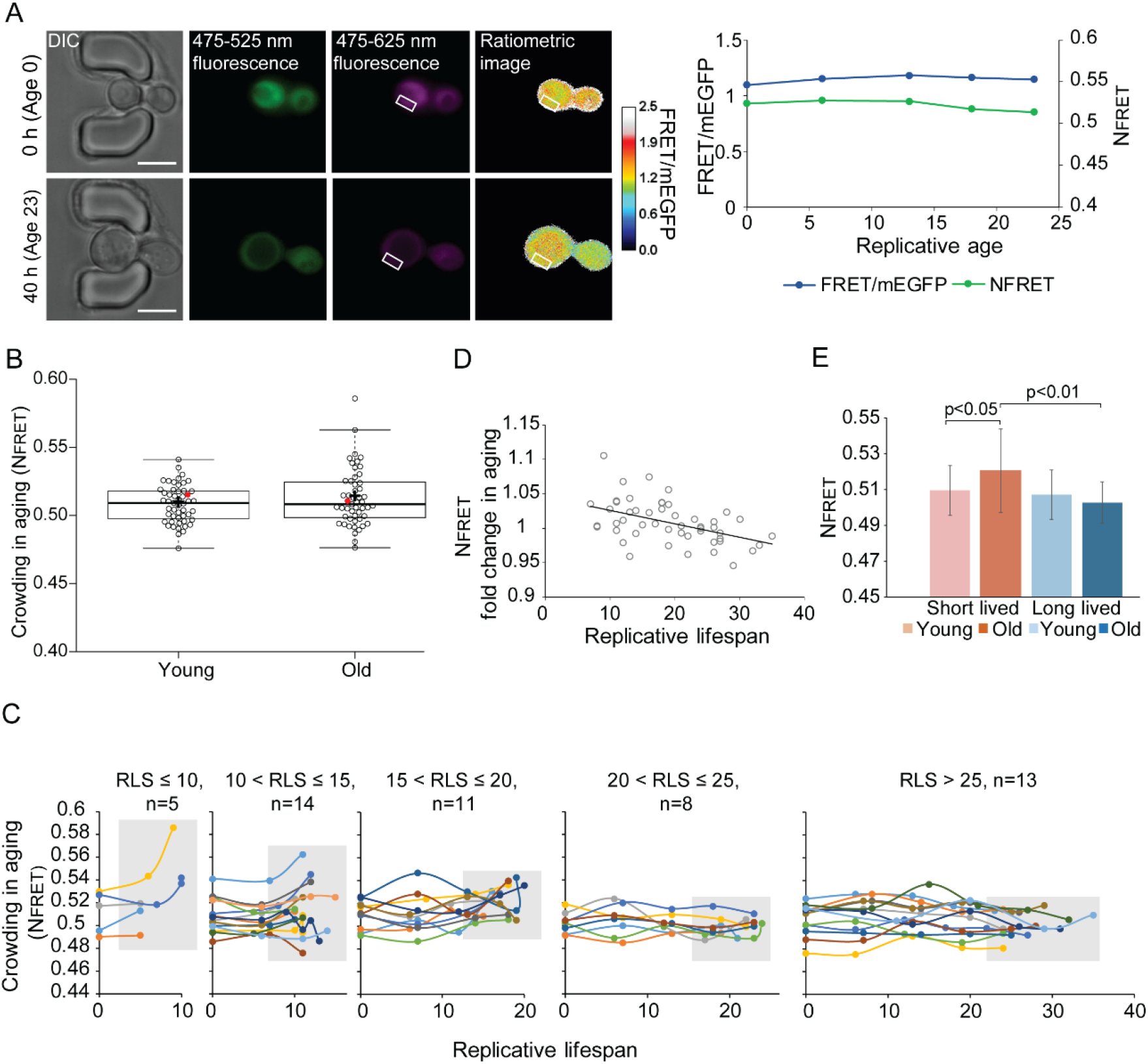
Crowding remains remarkably stable in aging. A) Left: Yeast cells expressing the crGE2.3 sensor, trapped in the aging chip. Images are from the same cell at the beginning of the experiment and the last measurement taken before cell death. Boxes indicate a cytoplasmic region. Right: Single-cell trajectory of FRET/mEGFP ratios (blue line) and normalized FRET (green line) B) Boxplot, showing normalized crowding ratios in young and old cells. For the young population, the first ratios recorded at the start of the experiment were taken. For the old population, the last ratio recorded before cell death was taken. The old population contains cells from different ages with a median lifespan of 19 divisions, n=51. The ratio recorded for the cell displayed in A is shown with a red dot. C) Single-cell trajectories of cells with indicated RLS-ranges and an indicated number of cells in each category. Grey boxes indicate the range of crowding ratios at the end of the RLS of each age group. D) The fold change in crowding plotted against the replicative lifespan of cells, R^2^ = 0.22, p = 5.9E-4; with Spearman correlation of - 0.4661, p = 5.7E-4) The average first and last recorded normalized crowding of cells with a RLS shorter (red) or longer (blue) than the median RLS of 19 divisions.

Plotting single-cell trajectories for cells that reach a replicative lifespan of 10, 10-15, 15-20, 20-25 or larger than 25 shows that the shortest-lived cells tend to increase the crowding levels during their lifespan, while the longer-lived cells tend to have more stable crowding levels (Fig. 3*C*). Indeed, there is a weak correlation (R^2^=0.13, p=0.009) between lifespan and old age crowding levels (Fig. S3*B*), and the fold change in N_FRET_ ratios in aging also shows a weak correlation with lifespan (R^2^=0.22, p<0.001) (Fig.3*D*). Also, in support of a relationship between crowding and aging, we observe that cells that live shorter than the median lifespan of 19 divisions have significantly higher ratios in aging (p<0.05), compared to long-lived cells (Fig. 3*E*). It seems that it is the maintenance of crowding homeostasis, rather than the absolute crowding levels, which has an association with lifespan, as lifespan does not correlate to the crowding ratios in young cells (Fig. S*3C*).

### Volume distribution of cellular compartments in yeast replicative aging

Cell volume increases in aging and the increase per division is predictive for the lifespan of cells^37^. Because macromolecular crowding is directly linked to cell volume, we aimed to determine how the volume of the cytosol changes in aging and assessed aged cells on the ultrastructural level. To specifically explore the ultrastructure of aged cells, we labelled the cell wall of cells in log-phase with Alexa-488, which were then allowed to age over 20 hours to achieve an estimated replicative age of 13 divisions. As the dye remains with the aging mother cells ^50^, we can specifically identify aged cells within the population using correlative light (Fig. 4*A*) and electron microscopy (Fig. 4*B*). 14 tomograms were acquired and segmented to define the plasma membrane, nucleus, vacuole, lipid droplets, ER, multivesicular bodies, and mitochondria (Fig. S4). The aged cells show diverse phenotypes, e.g. in terms of numbers and size of vacuoles, but in almost all cells, the vacuoles, take up a larger fraction of the cell volume than in young cells.

**Fig 4.**
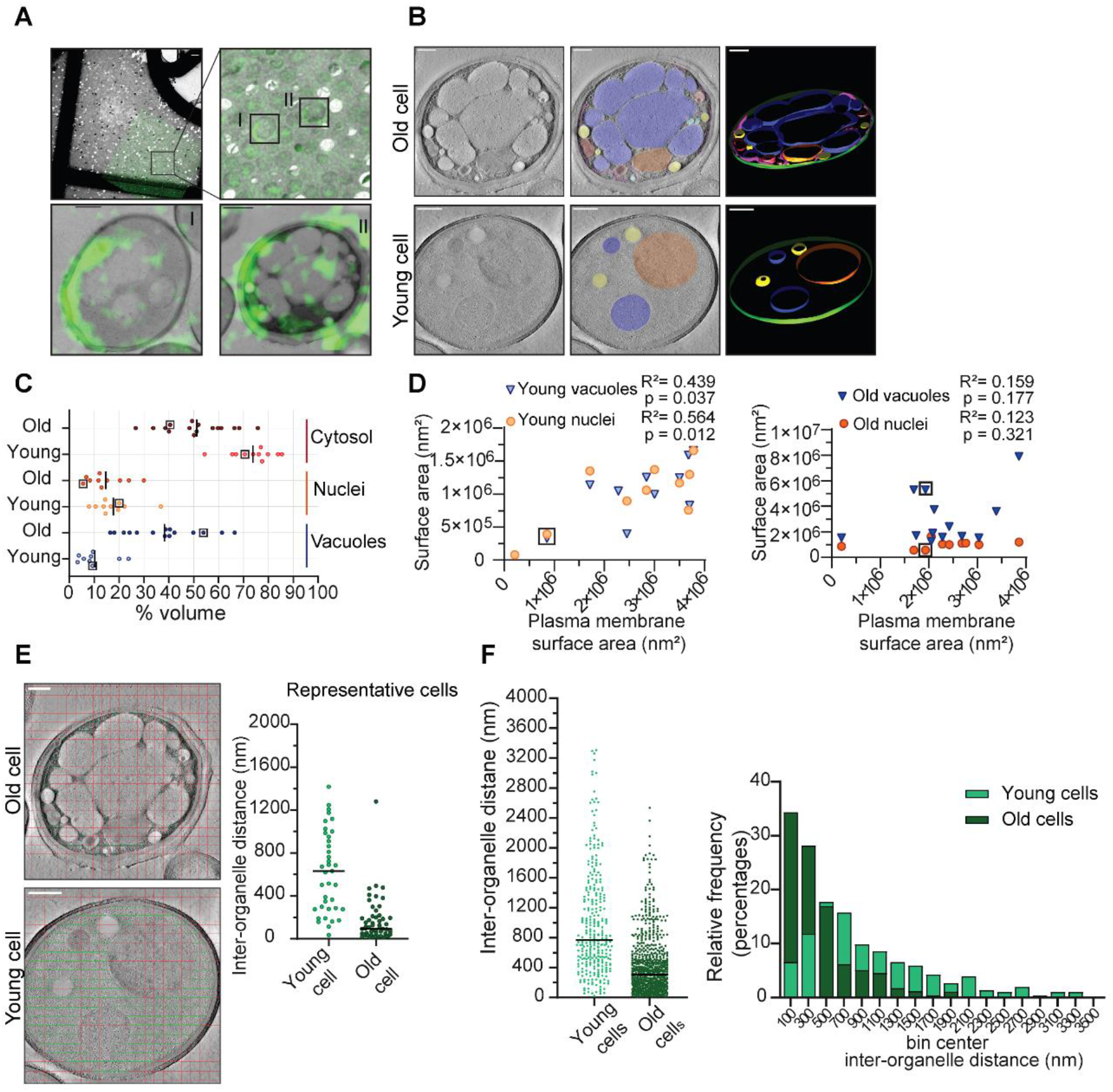
Inter-organellar crowding increases in aged cells. A) Identification of aged cells (20 hours, replicative age ∼13) by CLEM. Overlay of low magnification (225x) electron micrograph and fluorescence image and zoom-in of old cells with Alexa-488 labeling (boxed). Scale bar is 10 µm. Boxed cells I and II are shown at higher magnification (8900x) in lower panels with scale bar set at 1 µm. B) Representative single slices of tomograms without (left panel) and with an overlay to emphasize organelles (middle panel). 3D isosurface rendering (right panel) of tomograms of young or old cells. Nuclei (orange), vacuoles (blue), lipid droplets (yellow), mitochondria (red), ER (magenta), and plasma membrane (green). Scale bars are 500 nm. C) The relative cell volume occupied by vacuoles increases in old cells. The scatterplot shows volume fractions of nuclei and vacuoles in 14 aged cells and 10 young cells. The volume fraction of the cell section minus the volume fractions of the nucleus and vacuole is used to estimate of the cytosol volume fraction. The values for the example shown in B are boxed. Black lines denote the median. D) Membrane surface areas of nucleus and vacuole as a fraction of the plasma membrane surface area in young (left) and aged (right) cell sections. Each point represents a single nucleus or the sum of all vacuole surface areas within a single cell. The boxes indicate values from cells in B. R^2^ values were calculated from Pearson coefficient. E) Method to measure the distance between membranes in young and aged cells. Right is a scatter plot of measured distances from represented cells F) Left is a scatter plot of inter-organelle distances between membrane-bound organelles in young and old cells. The black line demarks the median. Right is a histogram of inter-organelle distance distribution. Values were grouped into 200 nm width bins. n=305 from 10 young cells, n=778 from 14 old cells.

To quantify organellar volume in the tomograms, we focused on the vacuole and nuclei, which are the largest compartments. We calculated their volume and membrane surface areas relative to the total cell volume and plasma membrane surface area in each tomogram (Fig. 4*C*). We find that the vacuoles take up 3 to 24% v/v in young cells and a higher volume fraction, namely 16% to 66% v/v, in old cells. The nuclei take up between 8 and 37% v/v in young cells, which is similar to old cells (7-30% v/v). Interestingly, we find that in old cells, vacuoles can occupy up to 66 % v/v, this is comparable to the theoretical maximum volume of 64% v/v that randomly packed spheres can occupy in a container.

Subtracting the nuclear and vacuolar volumes from the total volume provides an estimate of the cytosolic volume in the analysed tomograms. The cytosol volume takes up 54-82% v/v in the young-cell sections compared with 22-70 % v/v in those from old cells. Consistent with the changes in organelle volumes in aging, we find that the surface area of the vacuole and nuclear membranes relates to the surface area of the plasma membrane in the young cell sections. In contrast, the relation between organellar and plasma membrane surface areas is lost upon aging (Fig 4*D*). The loss of this relationship provides the correspondingly wide distribution in occupied organellar volumes.

To assess the consequences of the changes in organellar crowding, we determined the average distance between organelles (Fig. 4*E*), which would be a measure for cytoplasmic confinement of larger particles induced by organelles. We find a striking decrease in the most common distance from ∼500 nm in young cells to ∼100 nm in old cells, albeit with a wide distribution. Given a diffusion coefficient of 0.1 µm²/s previously determined in yeast cells for a particle with a diameter of 40 nm, it would take 0.005 sec versus 0.3 sec to reach a membrane with Brownian diffusion alone, assuming similar viscosity (supplementary information, Fig. S*5*).

Our analysis of the ultrastructure of aged cells further suggests that even though the total cell size increases in replicative aging cells ^7, 37–39^, the cytoplasmic volume fraction does not increase in aging but rather remains stable or even decreases as the vacuolar volume fraction increases. Moreover, we conclude that crowding at the length scales of organelles, which we coin organellar crowding, strongly increases with aging.

## Discussion

Here, we provide the first (to our knowledge) roadmap for the progressive decline in ageing in several parameters defining a cell’s intracellular environment, namely, pH, crowding, and volume, all of which impinge on the hallmarks of aging. We find that the largest changes arise from organellar crowding, while macromolecular crowding homeostasis is mostly maintained. The pH shows a progressive decline that follows the pI of the proteome. Below, we discuss possible causes of this changed intracellular environment and implications on molecular processes in an aging cell.

### pH homeostasis in aging

In general, pH has far-reaching consequences on cell physiology. Among others, the pH influences protein folding, enzyme activity, phosphorylation of metabolites and proteins, protein solubility/ phase separation, interactions between the molecules, redox potential, proton gradients, proton-dependent transport of nutrients ^11^. Even pH variations of 0.5 pH as we measure here, cause various enzymes to lose activity ^51, 52^ or induce liquid-liquid phase separations of proteins in cells ^53^.

The pH has been linked before to aging, showing that the vacuole and cell cortex get more basic^13^. Our results add to this work, showing that that pH is less well maintained in the cytoplasm during aging with overall acidification. The pH of the vacuole and cytosol become more similar in aged cells, which means that they lose an aspect of their compartment identity. Because the drop in cytosolic pH that occurs at later ages is not observed in all cells, and has limited predictive value for their lifespan, we can conclude that it is not a general early cause of aging, but instead could play secondary roles, such as driving entry into senescence and quiescence.

Can the decreasing pI of the proteome, observed in aging be a source for the drop in cytosolic pH? Cellular proteins have weakly acidic or basic residues, which collectively act as a buffer of cellular pH. It is proposed that yeast proteins evolved to have a pI that is similar to the pH of the compartment where they usually reside in ^54^, although this is not described in human cells ^55^. However, a cell must regulate its pH away from the pI to maintain protein solubility and prevent protein phase separation or aggregation. Thus, the yeast proteome may provide the basal pH of a cell compartment, while energy-dependent mechanisms regulate it away from the pI. This principle has been demonstrated *in vitro,* where isolated lysosomes maintain their acidity through Donnan-type equilibrium ^56^. Hence, if a cell’s energy state does not allow proper pH regulation, the pH would follow the pI. Therefore, the drop in pI of 7.1 to 6.6 could be responsible for the modest drop in pH during replicative aging, and when cellular energy is lost upon senescence, a more considerable drop ensues. In young cells, the vacuolar pH is around 5.5 ^57^, while its pI is 6.5 ^54^. To date, there are no quantitative data on vacuolar pH in aging, but it is known that the vacuoles lose acidity. It would be interesting to know whether pH levels in the vacuoles also approach values equal to the pI, as in the cytosol. Such observations raise the question of whether in old cells, the proteome becomes not only the main buffering mechanism, but also a regulatory mechanism for cellular pH. The cell-to-cell variation in pH that we observe during aging could, in turn, reflect variation in proteome composition or energy state. Additional mechanisms could add to the changes that we observe, such as altered proton transport activity, accumulation of acidic metabolites, or polynucleotide content. Nevertheless, the qualitative similarities between the pI and pH are striking.

### Crowding homeostasis in aging

Physicochemical parameters such as pH and macromolecular crowding can determine biochemical organization in yeast ^4, 19^ and potentially cause disease ^58^. Changes in physicochemical homeostasis could explain the multifactorial nature of the aging process since they will have ubiquitous and diverse effects on all cellular processes ^6^. We investigated macromolecular crowding as an intrinsic cell property, which can modulate e.g., phase separation and transition *in vivo* ^5, 19^. Considering the significant increase in volume the cell undergoes with age, it is at first somewhat surprising that we find crowding levels to be very stable in old yeast cells. This stability may be a consequence of a combination of relatively small changes in cytoplasmic volume fraction (discussed below) and efficient mechanisms to maintain macromolecular crowding homeostasis.

To date, the only way to change crowding levels in the cells considerably and rapidly is to apply an osmotic upshift in the medium. The cell responds immediately by uptake of potassium ions to regain its volume and crowding ^59^. Even in a potassium-deprived medium, the crowding effects are observed for a short period due to other response mechanisms. The short response time indicates that crowding levels (or thermodynamic water activity) are crucial. Crowding could be regulated by an array of mechanisms that prevent drift in crowding over more extended periods. These include e.g., i) uptake of counterions upon biopolymer synthesis, inducing an osmotic pressure over the membrane resulting in cell growth and reduction of the biopolymer concentration ^60^, ii) carbon catabolite repression to reduce the space taken up by metabolic enzymes ^61^, or iii) altering the ribosome/tRNA concentration ^5, 62^. These possible mechanisms suggest that crowding levels are tightly regulated in healthy cells ^34^, and we find here that this tight regulation expands to replicative aging.

Are there consequences of crowding during aging? We find that cells with slightly lower crowding levels that remain relatively unchanged during ageing have a longer lifespan, while cells increasing in crowding tend to be average or shorter-lived. Young cells display a natural variability of crowding between cells that is independent of lifespan. Therefore, retaining initial crowding levels could be beneficial for old cells. A too-large drop in crowding would reduce cell viability: It was recently suggested that dilution of the cytoplasm could evoke cell cycle arrest and lead to senescence, through a variety of mechanisms ^35^. Possibly, an optimum macromolecular crowding exists, from both a physicochemical and biochemical viewpoint, within a window of less optimal but viable crowding levels.

### Organellar size homeostasis in aging

Live-cell imaging studies of yeast expressing GFP-tagged organelle markers have highlighted how several organelles change in abundance or shape in aging yeast cells. Amongst them are the increase in vacuolar ^39^ and nuclear size, and fragmentation of nucleoli ^40, 41^ and mitochondria ^42^. However, because aged cells are scarce in exponentially growing cultures, the detailed ultrastructural properties of aged cells had not yet been researched using EM analysis. We use CLEM to reveal that aged cells have an altered ultrastructure. Particularly striking is that the available space for the cytoplasm can become minimal in aged cells and is enclosed by a large surface area of organellar membranes. The average membrane-to-membrane distance in aged cells is >2 times smaller in aged cells than in a young cell: The average distance between organelles decreases from ∼1000 to <500 nm. The distribution is, however, strongly tailing, and these averages correspond to a change in the most common distance from 400-600 to 0-200 nm. Already from the frequency distribution of the measured inter-organelle distances that are smaller than 80 nm, which is ∼12% in aged cells and ∼2% in young cells, one could deduce that contact sites are possibly expanded in aged cells. The implications of an increase in membrane contact sites can be widespread. Moreover, the enlarged compartmentalization must affect the movement of larger structures such as ribosomes and induce confinement on similar-sized particles in the 40 nm range.

The organellar crowding should have several effects that are highly dependent on the local distance between the membranes and the size of the particle, as demonstrated by the time required to diffuse to the membrane (supporting information, Fig. S*5*). The proximity to the membrane will also result in more interactions with membranes. The particles will also suffer an entropic cost by sacrificing translational degrees of freedom in the inter-organellar spaces. Hence, larger particles may be crowded out of regions with high organellar crowding leading to a size-dependent spatial sorting. On a technical note, given the extreme dependence on particle size, particles that are much smaller than the distance between the organelles would notice less of this confinement, which would include, for example, the macromolecular crowding sensor, which is polymer-like with a radius of ∼5 nm. In contrast, fluorescent particles with radii of ∼20 nm should experience confinement when present in the typical 100 nm confinements. Therefore, the behavior of such a particle will be dependent on where it is inside the cytoplasm. From a biological perspective, future studies should address how altered spatial sorting of large biological relevant particles, such as ribosomes, RNPs, or protein condensates or aggregates, would impact the cell physiology of aged cells.

### An integrated view of physicochemical homeostasis in aging

It is well established that aged cells are derailed in molecular aspects, such as the loss of protein homeostasis, genome stability or metabolic state, and that these molecular changes affect the cell’s physiology. However, the identity of cells and organelles is not only defined on a molecular composition level, but is also defined by physical and physicochemical properties. The data presented here highlights that these are also drifting in aging. First, the physical property of organelle size drifts in aging: where the cytosol in young cells occupies most volume, followed by the nucleus and vacuole (Fig. 4*D* *left*), in old cells the order is opposite: here the vacuoles are largest, and the cytosol represents the smallest volume fraction. In addition, the physicochemical property of pH changes in aging: where the pH of the vacuoles is kept much lower than the cytosol in young cells, their values come closer together as the cytosol acidifies and the vacuole loses acidity in aged cells ^13^. Lastly, crowding on the scale of organelles, which we term organellar crowding, sharply increases with aging. Organellar crowding likely influences phenomena that are on the 100 nm to µm length scale, such as long-range diffusion of larger particles and organelles, condensate formation, organellar shape, RNA translation, and cytoskeletal dynamics. Besides, the increased surface area presented by the organelles may give additional opportunity for adsorption or increased membrane contact sites. However, crowding at the length scale of a single protein, i.e., ∼10 nm, changes little, and these proteins do not experience a direct effect of organellar crowding. By analogy, an ant would not notice if there were a fence around a field, but an elephant does. We speculate that while in young cells, micrometer length structures are mostly hindered in their diffusion throughout the cytosol by cytoskeletal structures, in aged cells, the high occupancy of intracellular organellar membranes provides the major obstacle. In the context of these significant changes in the physical and physicochemical properties of aged cells, it is remarkable that macromolecular crowding on the scale of single proteins is rather stable in aging, and instigates future studies to identify what regulates these crowding levels.

## Materials and Methods

### Plasmid construction and yeast strains

All yeast strains (*Table S1*) were constructed in the BY4741 genetic background (*his3Δ1, leu2Δ0, met15Δ0, ura3Δ0*) ^63^ and were transformed with pRS303 yeast integrative plasmid, harboring the respective sensor gene with a *TEF1* promoter and *CYC1* terminator. A complete list of primers used to construct the strains can be found in Table S2.

To construct SMY008, the yeast codon-optimized gene of the crGE-NLS sequence was amplified from pYES2 vector (GeneArt, Invitrogen) together with pTEF1 and CYC1T by PCR with the forward primer F1_SM and reverse primer R1_SM, introducing SalI restriction site downstream of the terminator and removing the NLS localization signal. The sequence was subcloned into a pRS303 yeast integrative vector in between SpeI and SalI sites. The construct was then used to obtain chromosomal integration of the sensor sequence in the *HIS* locus.

For the generation of SMY015, the yeast codon-optimized mEGFP-crGE-mCherry (GeneArt, Invitrogen) was amplified using primer F3_SM to introduce a HindIII site and the primer R3_SM to introduce a stop codon and a downstream XbaI site. The PCR product was subcloned in pYES2-TEF1 between HindIII and XbaI. The resulting TEF1-mEGFP-crGE-mCherry construct was amplified with F1_SM an R1_SM and subcloned in pRS303, as described above. The gene encoding for Gamillus-crGE-mScarlet-I (GeneArt, Invitrogen) was amplified with primers F4_SM and R4_SM to introduce a stop codon, as well as XmaI and XbaI restriction sites after the mScarlet-I. The resulting PCR product was digested with NcoI and XmaI to isolate the mScarlet-I gene. The mCherry sequence in the pRS303-mEGFP-crGE-mCherry was then substituted with mScarlet-I, by subcloning between the NcoI and XmaI restriction sites. The resulting construct of mEGFP-crGE-mScarlet-I in pRS303 was used for chromosomal integration into the *HIS* locus of BY4741.

To construct SMY012, the pHluorin gene was amplified from pYES2-ACT1-pHluorin^64^ by PCR, using primers F2_SM, introducing HindIII restriction site and AAAAAA for enhanced expression in front of the start codon, and R2_SM, introducing a stop codon and XmaI and XbaI restriction sites. The PCR product was subcloned in the pYES2-TEF1 vector between HindIII and XbaI restriction sites. The TEF1-pHluorin-CYC1T construct was then amplified by PCR with primers F1_SM and R1_SM and integrated in the pRS303 with SpeI and SalI restriction sites. All pRS303 constructs were sequenced.

### Media and growth conditions

Yeast cells were grown at 30 °C, 200 rpm in Synthetic Dropout medium without histidine (SD-his), supplemented with 2% (w/v) glucose. Cells from an overnight culture are diluted 100× in 10 mL of SD-his, 2% glucose. After 7 h of incubation, appropriate dilutions were made to obtain cultures in the exponential growth phase on the following day (OD_600_ = 0.4−0.7).

### Microscopy

All *in vivo* experiments were performed at 30 °C. Images were acquired using a DeltaVision Elite imaging system (Applied Precision (GE), Issaquah, WA, USA) composed of an inverted microscope (IX-71; Olympus) equipped with a UPlanSApo 100× (1.4 NA) oil immersion objective, InsightSSI solid-state illumination, ultimate focus, and a PCO sCMOS camera.

Excitation and emission were measured with the following filter sets, respectively, and in the indicated order: crGE: CFP 438/24 and 475/24 nm, YFP 513/17 and 548/22 nm, FRET 438/24 nm and 548/22 nm. crGE2.3: FICT: 475/28 and 525/48 nm, A594: 575/25 and 625/45 nm, FRET: 475/28 and 625/45. pHluorin: DAPI: 390/18 and 435/48 nm, FITC: 475/28 and 525/48 nm. For crGE and pHluorin, 32% transmission power and for crGE2.3 10% for the FITC channel and 100% for the A595 and the FRET channel. For the aging experiments, stacks of 3 or 4 images with 0.7 μm spacing were taken, and for other experiments, stacks of 30 images with 0.2 μm spacing were taken at an exposure time of 25 ms for all experiments.

### Image analysis

Processing of all images was performed using Fiji (ImageJ, National Institutes of Health). For each image, the z-stack with the best focus was selected. pHluorin and the crowding sensors localize in the cytosol and nucleus, and appear to be excluded from the vacuole and probably also from other membrane-enclosed cytoplasmic organelles. We determined the fluorescence in each channel for the entire cell and subtracted the background from a region outside the cell. The respective ratios were subsequently calculated.

### Relation between pHluorin ratios and pH values

As described in ^4^, 2 mL of exponentially growing culture with OD_600_ of 0.5 were centrifuged at room temperature for 3 minutes at 3,000g in a tabletop centrifuge. The cells were then resuspended in 200 μL calibration buffer (50 mM MES, 50 mM HEPES, 50 mM KCl, 50 mM NaCl, 200 mM NH_4_CH_3_CO_2_) at pH 5, 5.5, 6, 6.5, 7, 7.5, and 8. We added 75 μM monensin, 10 μM nigericin, 10 mM 2-deoxyglucose, and 10 mM NaN_3_ (final concentrations) to each buffer. The cells were then loaded in microfluidic chips (see below), and their fluorescence was determined (Fig 1B).

### Aging experiments

The microfluidic chips were used as described previously ^7^. DIC images of the cells were taken every 20 min to follow the number of divisions for each cell and determine the replicative lifespan. Fluorescent images were collected every 10 hours. Z-sections were taken in both DIC and fluorescence imaging with 3 or 4 slices of 0.7 μm thickness. The experiment was left to continue up to 80 h and only cells that were trapped from the beginning of the experiment and died in the device within the time course of the experiment were included in the analysis.

### pH sensitivity of crGE and crGE2.3 crowding sensors

Cells from exponentially growing cultures at OD_600_ of 0.5 were harvested and resuspended in the same calibration buffers titrated to pH of 5, 5.5, 6, 6.5, 6.7, 7, 7.3, 7.5, 8 as the pHluorin calibration. The FRET/CFP and FRET/mEGFP ratios were determined from cells on a glass slide.

### Cycloheximide treatment

A 20 mL exponentially growing culture was split into two cultures of 10 mL. To one of the flasks, a final concentration 1 μM cycloheximide was added from a 1000× stock solution in DMSO. As a control, 10 μL DMSO was added to the other culture. Samples were collected immediately after the addition of cycloheximide or DMSO and imaged. Both cultures were then incubated at 30°C, shaking at 200 rpm for 90 minutes. Samples were collected after 90 minutes from both treatment and control cultures and imaged as described before.

### Osmotic shock

2 mL of exponentially growing culture was collected by centrifugation at 3,000 g for 3 min. The cells were resuspended in 200 μL low osmolality buffer (50 mM NaPi, pH 7 for isotonic conditions) or high osmolality buffer (50 mM NaPi, 1 M NaCl or 1.5 M Sorbitol) to induce the osmotic upshift. Cells were then placed immediately on the glass slide and imaged.

### Determining N_FRET_

Fluorescence signals from the donor (I_mEGFP_), acceptor (I_mScarletI_), and FRET (I_FRET_) channels after background subtraction were used to calculate the normalized FRET (N_FRET_)^49^. We did not correct for the donor bleedthrough in the FRET channel and the acceptor cross-excitation because these contributions were minimal.

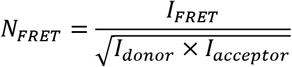

### Proteome isoelectric point

To calculate the overall isoelectric point of the aging proteome, we used published datasets for the age-related change in protein abundance ^14^, the protein copy numbers in young cells ^47^ and the computed isoelectric points (pI) (Saccharomyces Genome Database, SGD, https://www.yeastgenome.org/). We excluded 264 proteins from the aging dataset for which a copy number was not available. Out of the remaining 1229 proteins, 1071 proteins belong to the GO term ‘cytoplasmic component’ (Panther Gene Ontology; www.pantherdb.org), indicating the aging proteome mostly reflects (highly abundant) cytosolic proteins. For each time point in aging, the contribution of and individual protein to the overall pI was calculated by multiplying its pI with its relative abundance in the proteome. The total proteome pI was then derived as a sum of all weighed isoelectric points of the proteins in the dataset.

### Ultrastructural analysis of aged cells

WT cells were grown to mid log phase in SD medium supplemented with 1% glucose. 8 x 10^7^ cells were collected by centrifugation, washed twice with PBS and resuspended in 0.5 mL of PBS. 4 mg of EZ-link Sulfo-NHS-LC-Biotin (ThermoFisher Sci.) was dissolved in ice-cold H_2_O and immediately mixed with the cell slurry, which was incubated for 20 minutes at RT. Biotin-labeled cells were collected by centrifugation and washed twice with PBS. After resuspension in 100 μL of PBS, 5 μL of 5 mg/mL streptavidin-labeled with Alexa Fluor 488 (ThermoFisher Sci.) was added for 30 minutes. After washing in PBS, 40,000 cells were used to inoculate 30 mL of SD with 1% glucose, which was grown to an average age of 13 before processing for CLEM as follows. Cultures were concentrated into a thick slurry by gentle centrifugation and aspiration of excess media. These samples were high-pressure frozen in a Leica HMP100, freeze-substituted in a Leica Freeze AFS with 0.1% uranyl acetate in dry acetone, and infiltrated with Lowicryl HM20 resin. The polymerized resin block was cut into ∼200 nm thick sections onto 135 mesh H15 patterned copper/rhodium grids (Labtech). Fluorescence imaging was carried out as previously described in ^65^. Fluorescent micrographs were acquired using a DeltaVision widefield microscope (Applied Precision/GE Healthcare) equipped with UPlanSApo 60x (1.64 NA) and 100x (1.4 NA) oil immersion objectives (Olympus), solid-state illumination and CoolSnapHQ^2^ CCD camera (Photometrics). Bright-field and fluorescent images of grid squares with cells were acquired at both 60x and 100x magnification to facilitate alignment to electron micrographs in subsequent steps.

Grids were post-stained with lead citrate and labelled on both faces with 15 nm gold fiducials. Tilt series from −60° to 60° of selected cells were acquired with a magnification of either 8900x or 13300x on a FEI F20 fitted with a FEI Eagle CCD camera (4k x 4k) and using Serial EM ^66^. Tomograms were reconstructed in IMOD ^67^ using 15 nm gold fiducials for alignment. Low magnification (225x-440x) electron micrographs were also acquired to facilitate alignment with fluorescence micrographs and tomograms.

Correlation between light micrographs and electron micrographs was completed in Icy ^68^ using the ec-CLEM plugin ^69^. Corresponding points in EM and light microscopy images were selected based on cellular features distinguishable in both the light microscopy and EM.

### Analysis of electron tomograms

Manual segmentation of tomograms was performed in IMOD/3DMOD (Version 4.9.8,^67^) with contours drawn every 30 nm or less in z. The surface area and organelle volume were calculated from uncapped meshed models in IMOD. The cytosolic volume was estimated by subtracting the calculated volume of nuclei and vacuoles from the volume encapsulated by the plasma membrane.

Inter-organelle distances were calculated in an unbiased manner by overlaying horizontal lines spaced every 200 nm over the midplane of the reconstructed tomogram using the stereology tool in IMOD. Distances between organelles on overlaid lines were then drawn as contours and measured. As cortical ER was not faithfully visible in each cell, all ER membranes were excluded from the analysis. Graphs were compiled in Prism (GraphPad). The linear correlation coefficients (Pearson coefficient, r) and R^2^ were calculated in Prism (GraphPad).

## Acknowledgments

We acknowledge The Netherlands Organization for Scientific Research: Vidi grant 723.015.002 to A.J.B. and BBoL grant 737.016.016 to L.M.V., and the NIH RO1 GM105672 to D.J.T. and C.P.L., and R01 AG056359, and P30 AG013280 to MK for financial support.

## Supporting information

**Table S1.**
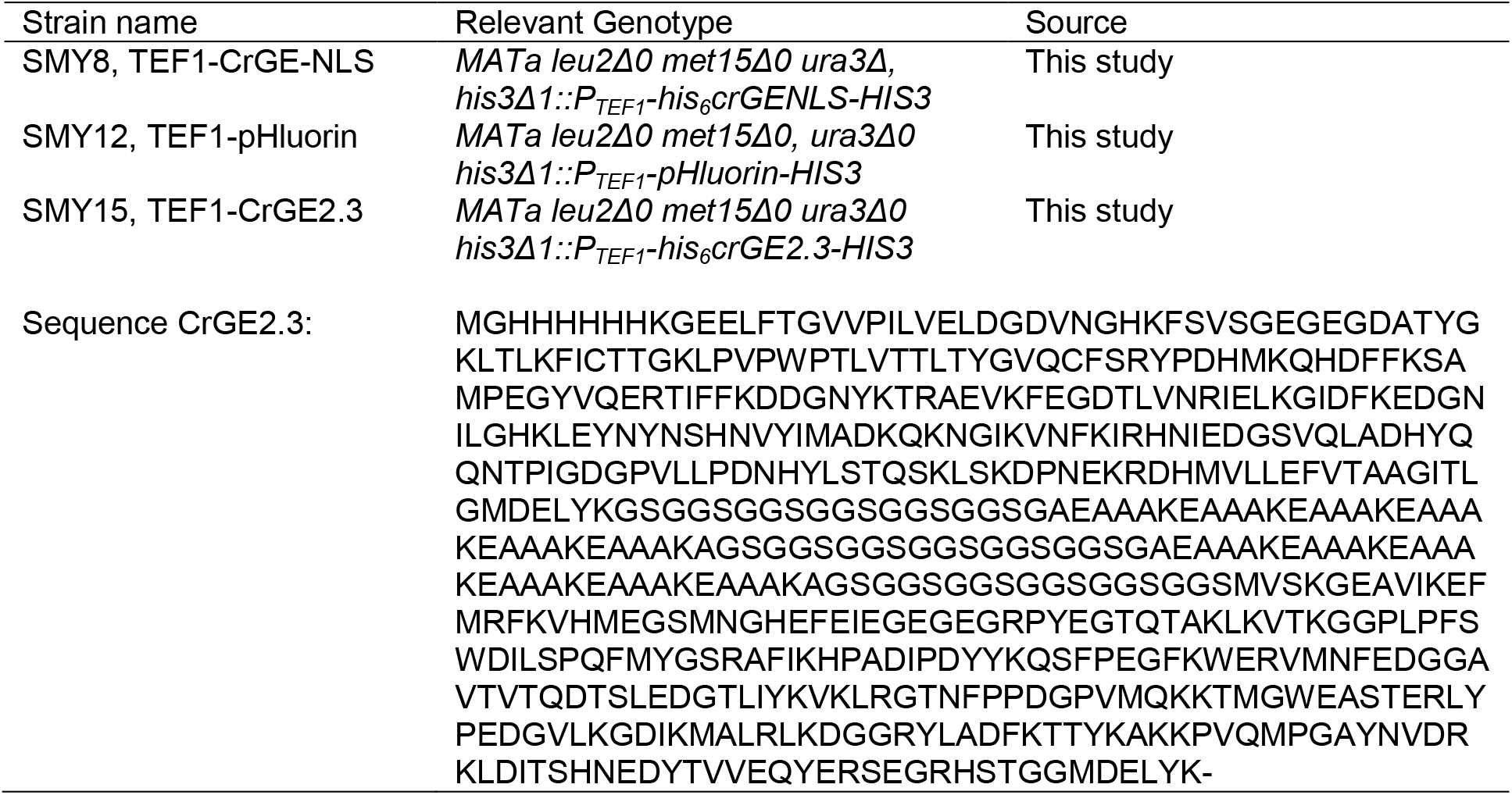
Strains used in this study.

**Table S2.**
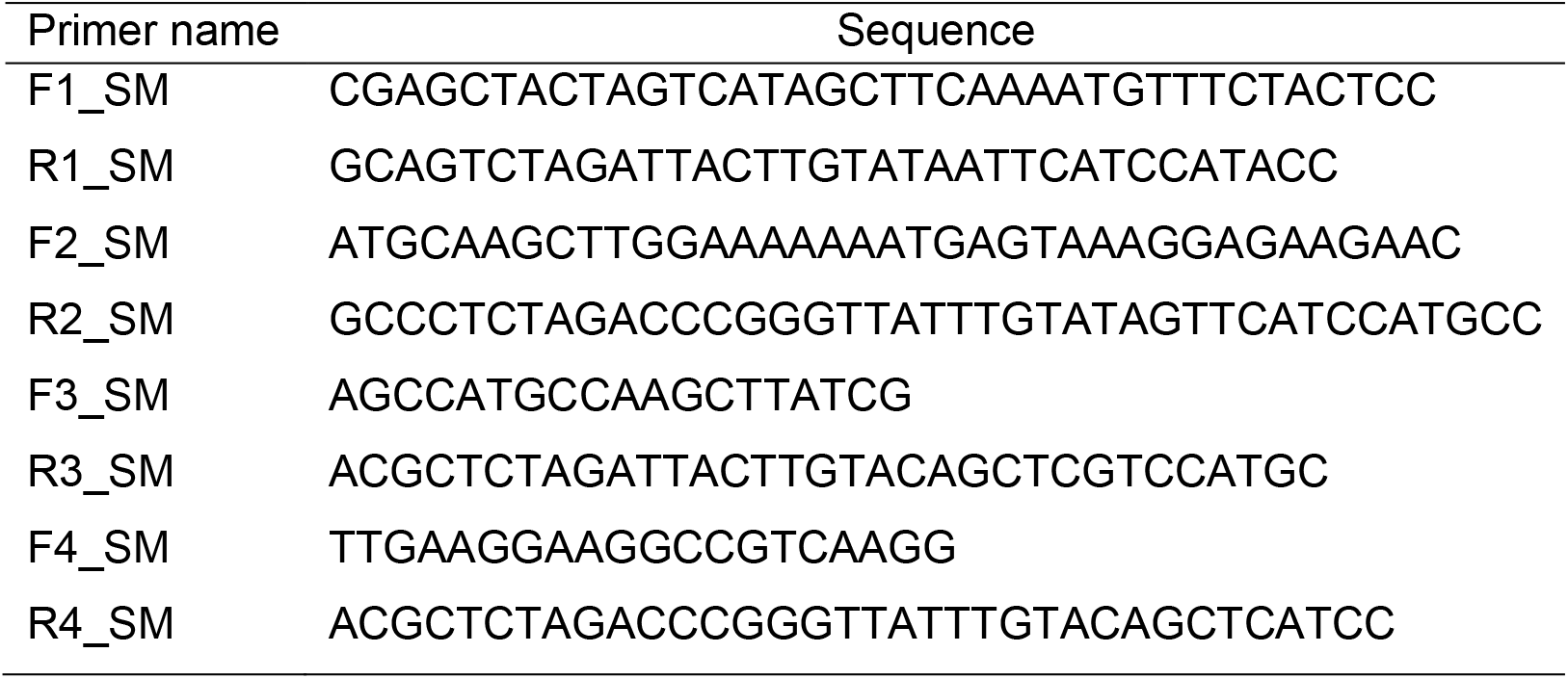
Primer sequences used in this study.

## Figures and Tables

**Figure. S1.**
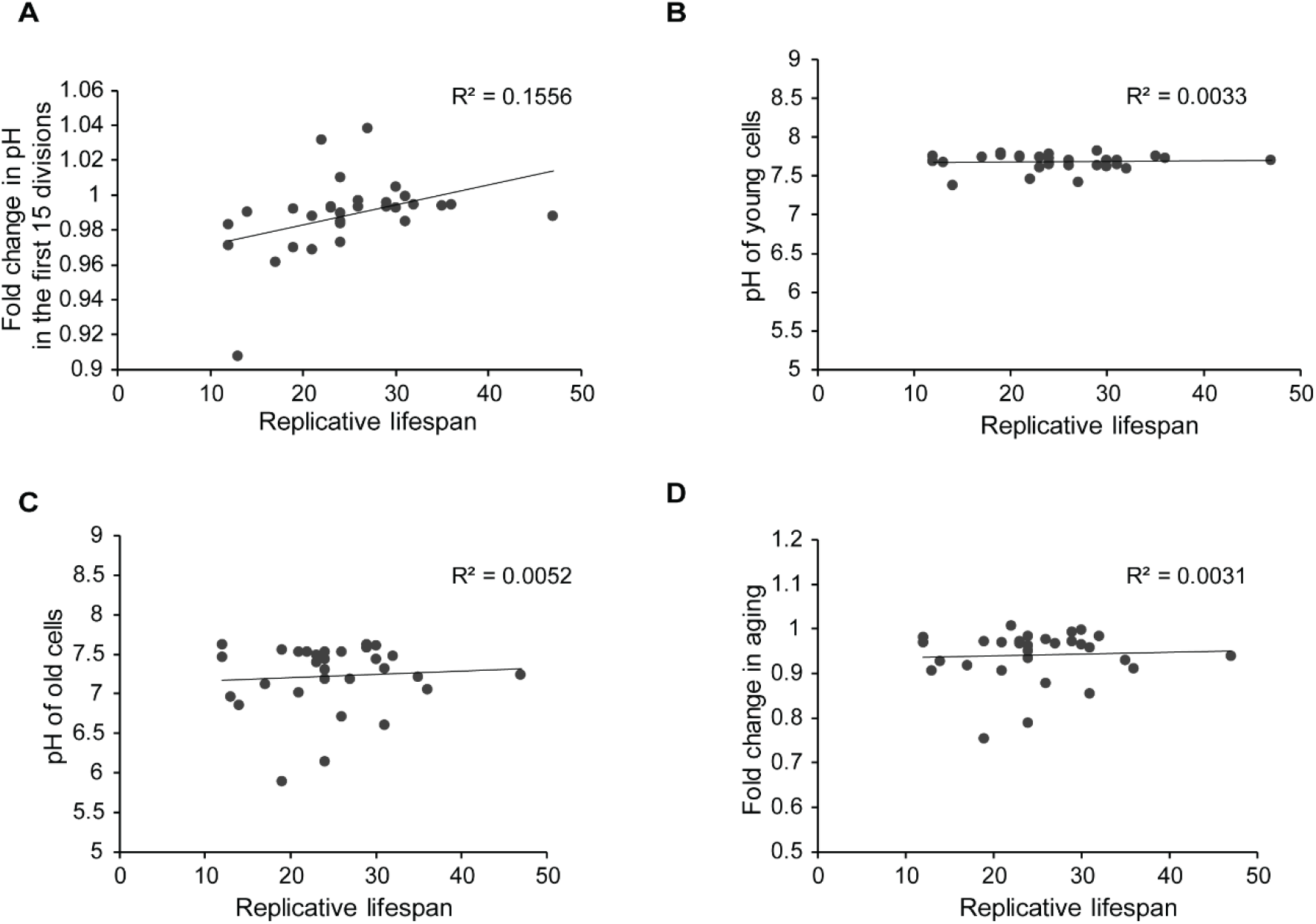
Age-related changes in cytosolic pH are not predictive for lifespan. (A) The change in pH within the first 15 divisions correlates weakly with lifespan, R² = 0.1556, p = 0.031006, shown also by Spearman correlation (r_s_) of 0.5564, p = 0.0014. However, the pH in (*B*) young, r_s_ = −0.1466, p = 0.4394 and (*C*) old cells, r_s_ = 0.038, p = 0.9841, as well as the (*D*) pH fold change, r_s_ = 0.042, p = 0.8258, do not correlate with the replicative lifespan.

**Figure. S2.**
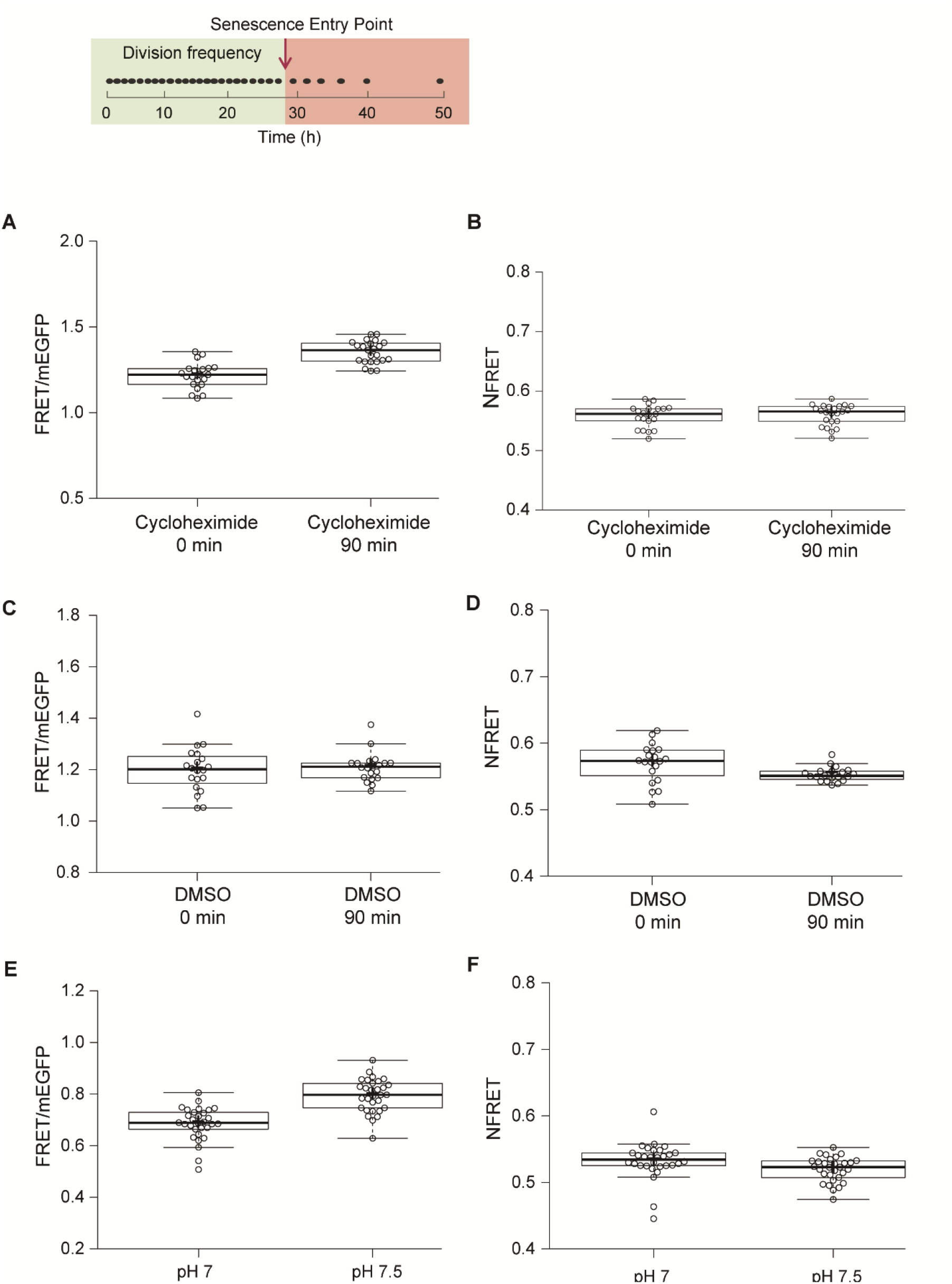
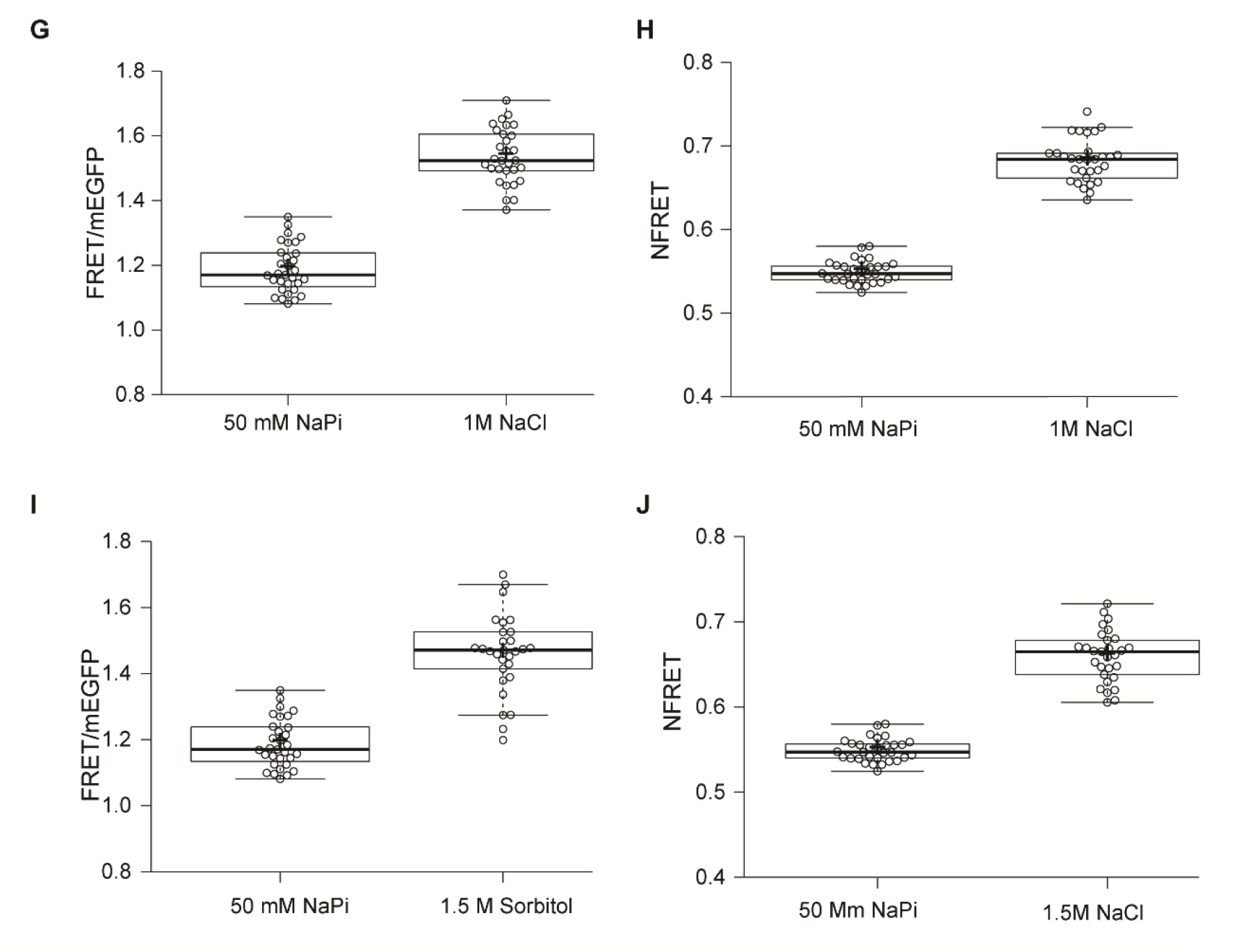
Readouts from fluorescent sensors are sensitive to conditions affecting the amount of actively fluorescent donor and acceptor of FRET, such as changes in division frequency and varying intracellular pH. (*Top row*) The division frequency of a representative cell during its replicative lifespan in the microfluidic device, indicated by black dots. The timeline is indicated underneath. The green box highlights the part of the lifespan of the cell, where the division frequency is constant. The supposed senescence entry point is indicated with an arrow, and the red box highlights the reduced division frequency after the cell enters into senescence mode. (*A*) Cells from an exponential culture, expressing the new version of the crowding sensor CrGE2.3, are treated with cycloheximide or (*C*) DMSO as control, and samples were collected at two points: 0 minutes and 90 minutes after the addition of cycloheximide or DMSO. FRET/mEGFP ratios increase in the cycloheximide probe, but not in the DMSO probe, showing the sensitivity of the sensor to the different maturation kinetics of the donor and acceptor. (*B*) When applying a correction for an unequal number of donor and acceptor, as described in the methods section, there is no difference in the normalized readout after 90 minutes of cycloheximide treatment and (*D*) a small reduction in DMSO treatment. (*E*) The sensor is mildly sensitive to pH changes within the relevant physiological range. (*F*) When we normalize the readout, this dependence is alleviated. (*G*) The crowding readout reports on increased crowding through hyperosmotic shock, induced by 1M NaCl or (*I*) 1.5 M Sorbitol. (*H and J*) The normalization procedure does not affect the increase in crowding readout, induced by actual crowding events. To test if the correction compensates for maturation artifacts, we subjected cells to 90 minutes of cycloheximide treatment. Cycloheximide inhibits protein translation and allows the fluorescent proteins to fully mature. We compared the FRET ratios before and after 90 minutes of treatment with 1 μM cycloheximide and estimated an 11% increase with average ratios starting from 1.22 immediately after addition of cycloheximide to 1.35 after 90 minutes of treatment (Fig. *S2A*). We added 10 μL DMSO to the control culture and did not observe any significant changes in the average ratio, which remains 1.2 after 90 minutes of exposure to DMSO (Fig. *S2C*). The increase in the FRET/donor ratio observed upon cycloheximide treatment is likely related to maturation instead of crowding because inhibiting protein synthesis would not increase crowding. The N_FRET_ ratio did indeed not change during cycloheximide treatment, showing the validity of correction for an unequal number of donor and acceptor due to maturation artifacts (Fig. *S2B*). We observed similar effects for the pH: Permeabilized cells at pH 7 and pH 7.5, which is a physiologically relevant pH range in the context of ageing and a range where the mScarlet-I is pH sensitive, gave FRET ratios at pH 7 of 0.69 and 0.79 at pH 7.5 (Fig. *S2E*), while the corresponding N_FRET_ ratios are the same with 0.53 and 0.52, respectively (Fig. *S2F*). Importantly, this normalization retains readouts from crowding changes (Fig. *S2*), where the increase remains 20% upon osmotic upshift (Fig. *S2G-J*).

**Fig. S3.**
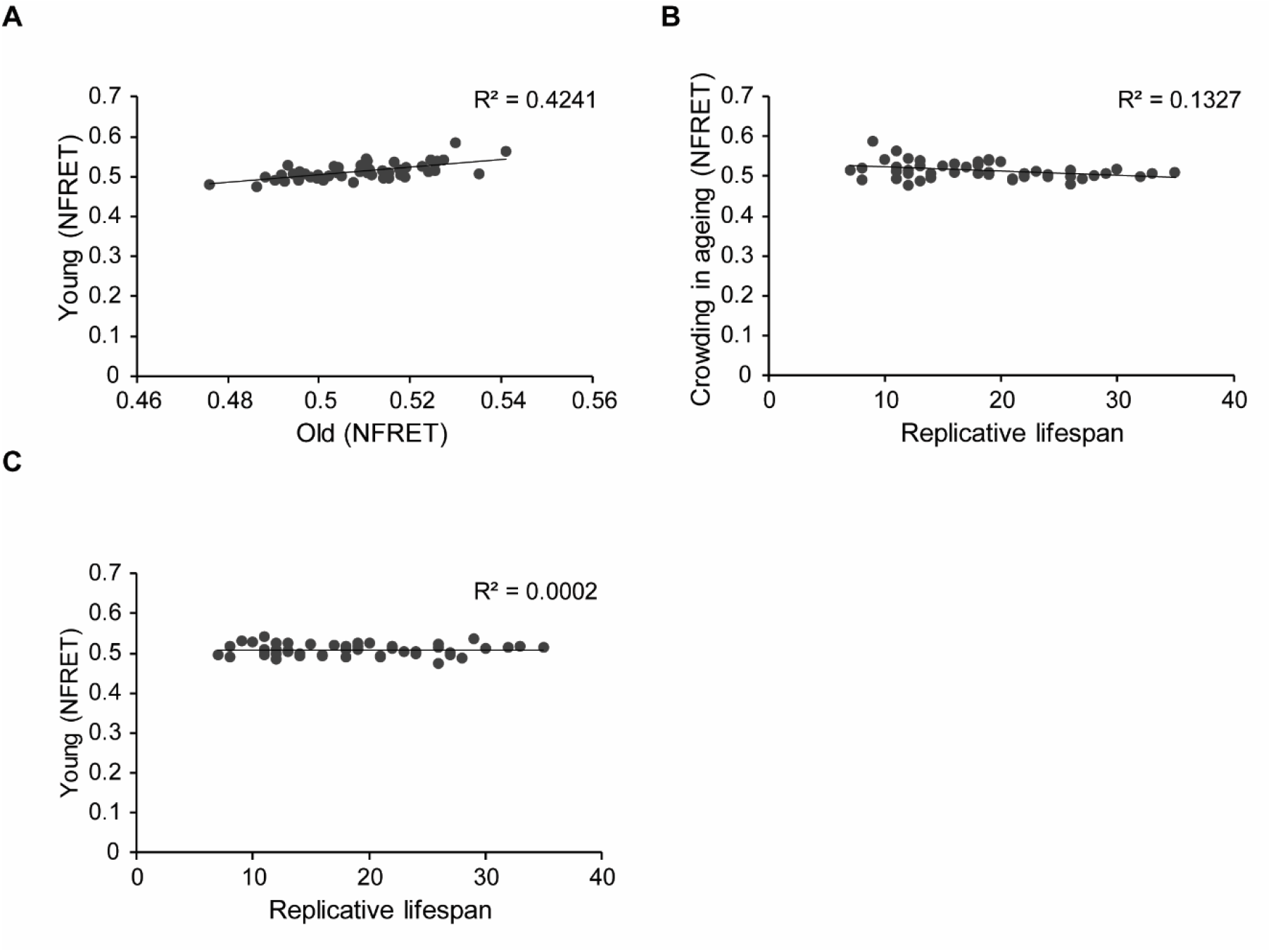
Crowding levels vary between individual young cells, but starting crowding levels do not associate with lifespan. However, in old cells, there is a weak correlation between lower crowding levels and longer lifespan. (*A*) Normalized crowding readouts from young cells, plotted versus old cells, show that crowding levels vary between individual cells and are generally maintained throughout aging; R^2^=0.4241, p = 2.27949E-07, also shown by Spearman correlation of 0.6239, p = 1.82E-06. (*B*) There is a weak correlation between crowding levels in old cells and lifespan, R^2^ = 0.1327, p = 0.00858559, r_s_ = −0.3518, p = 0.0114. (*C*) Starting crowding levels are not predictive for lifespan, shown by the lack of correlation between crowding readout in young cells and their lifespan, r_s_ = −0.0034, p = 0.9814.

**Fig. S4.**
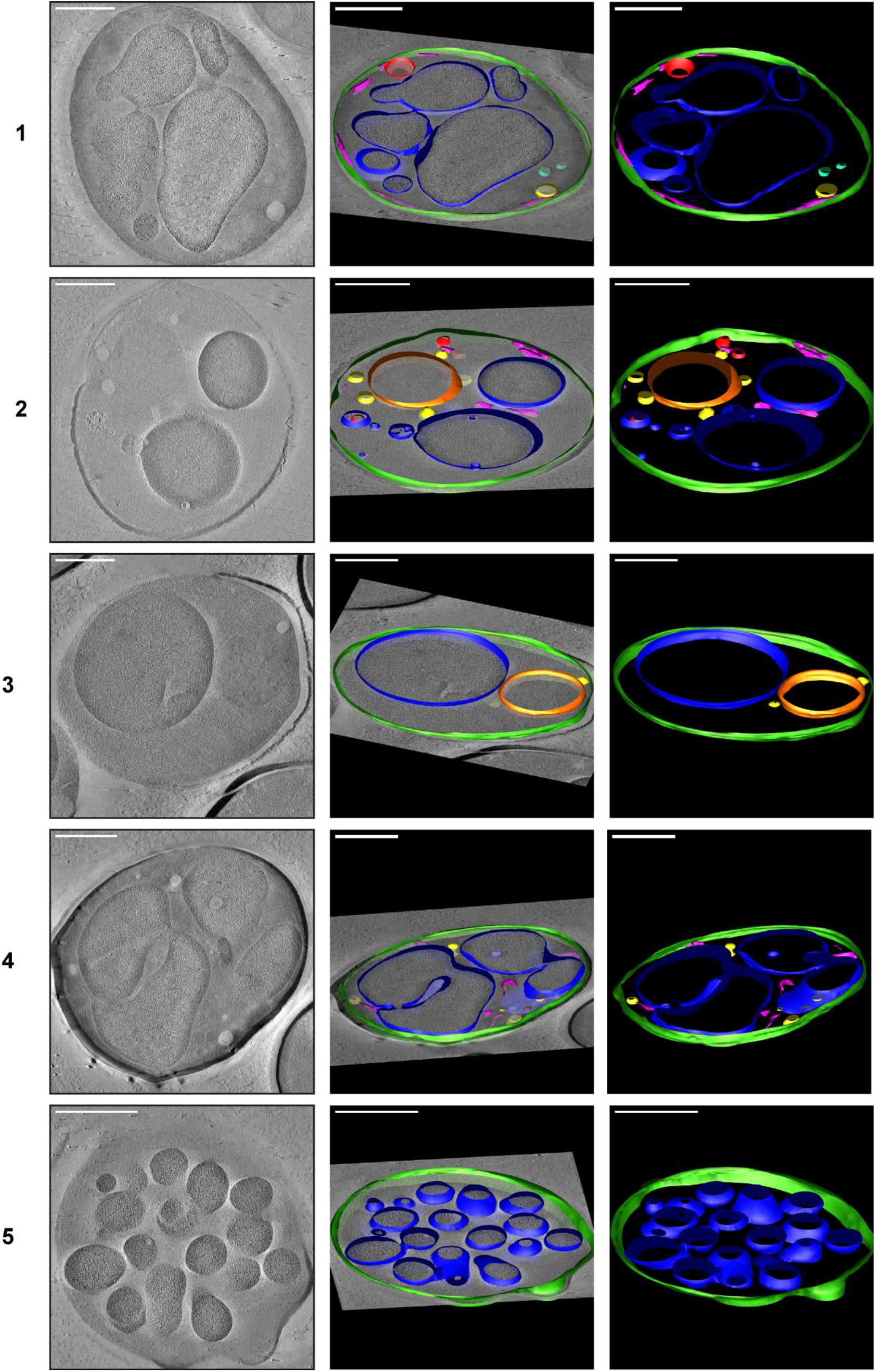
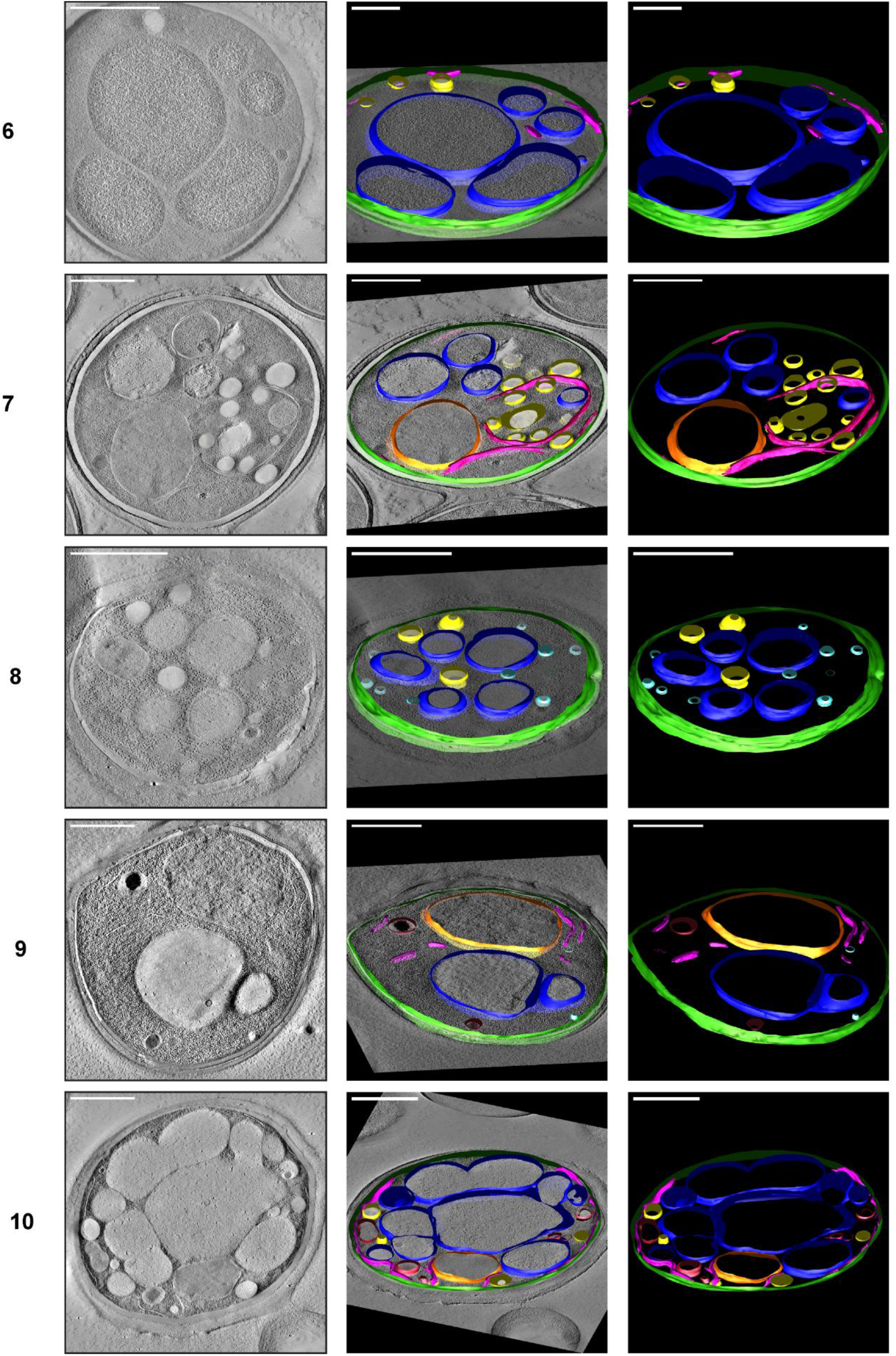
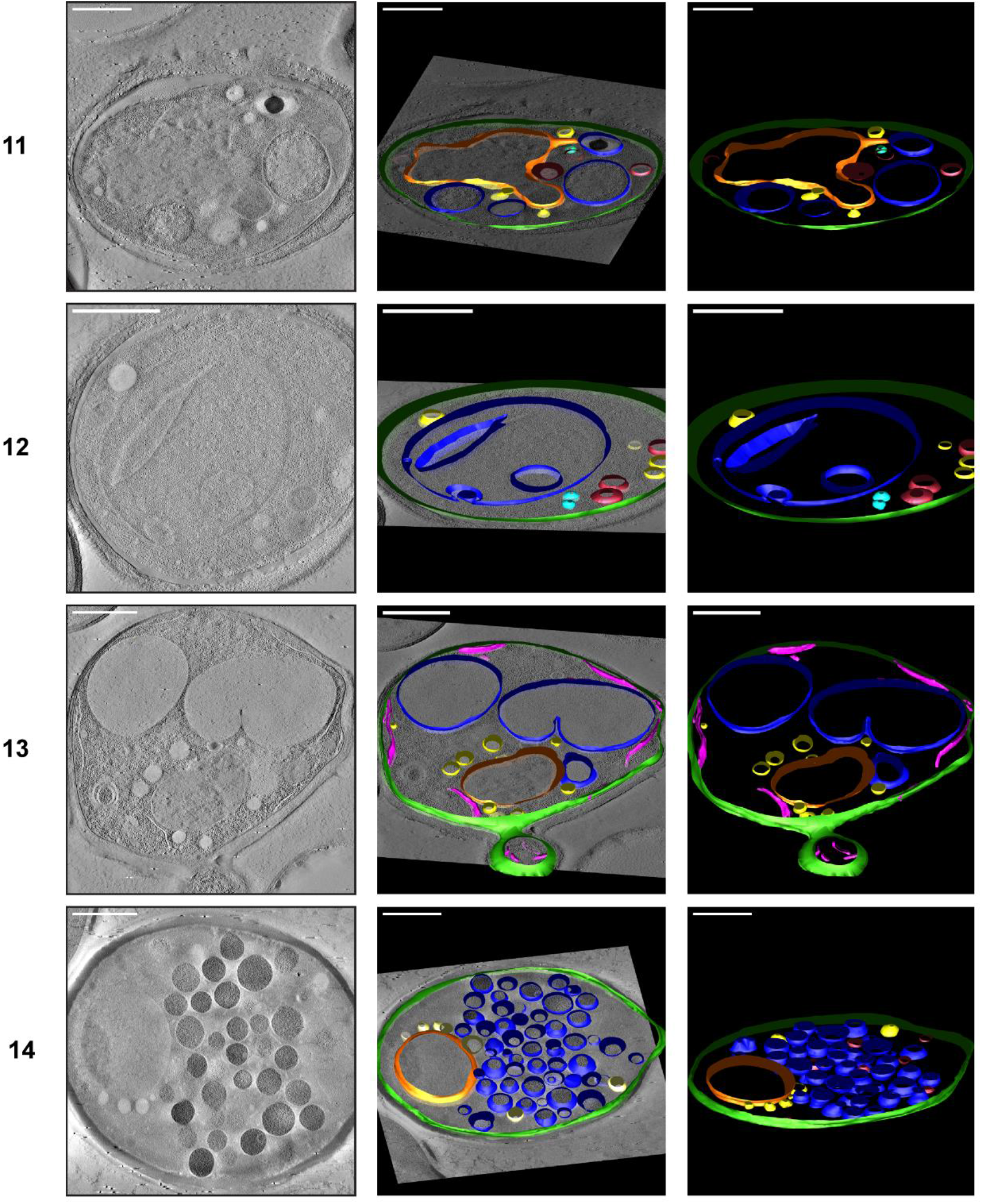
3D isosurface rendering of tomograms of 14 aged cells (around age 13). The sections show that plasma membrane (green), nucleus (orange), vacuole (blue), lipid droplets (yellow), ER (magenta), and mitochondria (red) can be detected. The aged cells show diverse phenotypes, but in most cells, the organelles, particularly the vacuoles, take up a much more significant fraction of the cell volume than in young cells. Scale bars are 1 μm.

**Figure S5:**
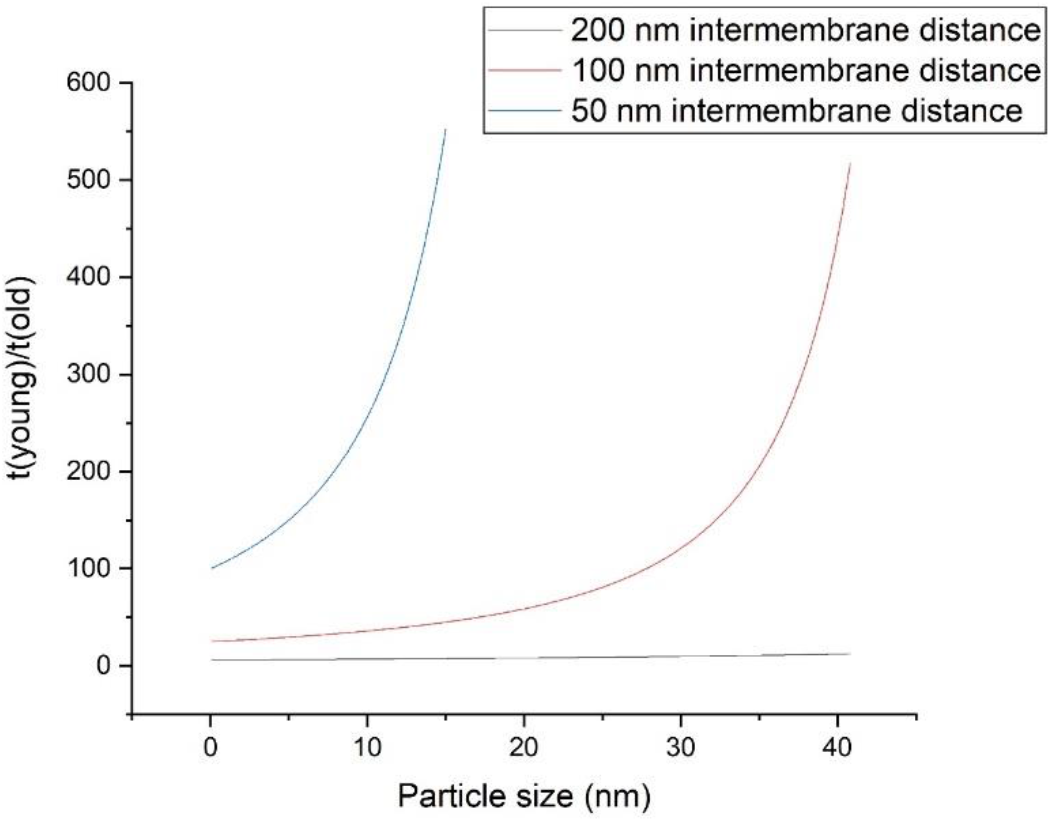
Strong particle-size dependence on the time to diffuse to the membrane. The ratio is independent of the diffusion coefficient, assuming that old and young cells have similar viscosity. The intermembrane distance of young cells is set at 500 nm, and the red line is a distance of 50 nm, green 100 nm, and blue 200 nm in old cells. 100 nm distances are most frequently observed in aged cells albeit with a wide distribution in distances. Note that in this analysis, a 20 nm particle would already result in 60 times shorter diffusion time to a membrane compared to a young cell, while a 40 nm particle results in 440 times shorter diffusion time.

## Dependence of particle size on diffusion time to a membrane

The weighted mean distance *d* a particle travels by Brownian diffusion depends on the diffusion coefficient *D* and the time *t* following the equation ^70^:

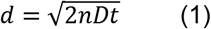

Here *n* is the dimensionality, which we set to 1. In reality, the particle can move in 3 dimensions. We set *d* as half the distance between the organelles, which is the distance the particle has to travel to the walls. The resulting *t* provides the time a particle diffuses on average to hit the wall. We thus assume the particle starts in the middle, and therefore the time estimates are longer than when it would start at another point.

This formula is derived from the chance *c* a particle has traveled to position *x*, which is at the organelle:

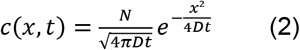

The chance can be set arbitrarily to determine the time it requires for a particle to interact with the organelle. This formula will result in the same trends as equation (1) for observing the relative effect of the change in organelle distance upon aging. We, therefore, continue with equation (1).

We take the distance the particle has to travel as *d=L/2*, with *L* the distance between the organelles. To take into account the size of the diffusing particle, we set *d=L/2-r*, with *r* = radius of the particle. Inserting into equation (1) gives for the average time for a particle to diffuse to the organelle:

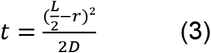

To compare old cells with young cells, we take the ratio *t*(young)/*t*(old) that provides the fold-decrease in time required to diffuse to the organelle membrane.

